# Transcriptomic entropy reveals tissue-specific patterns in aging and predicts cancer progression

**DOI:** 10.1101/2025.06.29.662218

**Authors:** Gabriel Arantes dos Santos, José Pedro Castro, Pedro A F Galante

**Affiliations:** Hospital Sírio-Libanês, São Paulo, SP, Brazil; i3S, Instituto de Investigação e Inovação em Saúde, Universidade do Porto, Porto, Portugal; Aging and Aneuploidy Laboratory, Instituto de Biologia Molecular e Celular, Universidade do Porto, Porto, Portugal

**Keywords:** Functional genomics, Geriatric oncology, Biogenrontology

## Abstract

Aging and cancer share complex molecular mechanisms, yet distinguishing between causative factors and byproducts remains challenging. Here, we investigated the role of transcriptomic entropy in aging and cancer processes by analyzing RNA-sequencing data from thousands of human and mouse samples. We found that entropy changes during aging are highly tissue-specific, with some tissues showing increased entropy while others exhibit decreased or stable entropy levels. Transcriptomic entropy strongly correlates with age-related processes, showing positive associations with proliferation, cellular senescence, and somatic mutation burden, while negatively correlating with stemness. Surprisingly, cellular reprogramming also increases transcriptomic entropy. In cancer, we observed that primary tumors generally display higher entropy than normal tissue, with entropy levels further increasing in metastatic stages. Notably, treatment-resistant tumors showed distinct entropy patterns, with acquired resistance associated with increased entropy, while primary resistance and immediate post-treatment responses showed decreased entropy. Higher entropy levels predicted poor survival outcomes in multiple cancer types, suggesting its potential as a prognostic marker. Furthermore, differential expression analysis revealed that entropy-associated genes are enriched in developmental processes and depleted in metabolic pathways, indicating a possible link to cellular dedifferentiation. Finally, we found increased entropy in various age-related diseases beyond cancer, suggesting that transcriptomic disorder may be a common feature in age-related pathologies. Our findings establish transcriptomic entropy as a fundamental parameter in aging and cancer progression, offering new insights into disease mechanisms and challenging the current view that increased disorder is always detrimental.

## INTRODUCTION

Despite significant advances in aging biology, distinguishing between causative mechanisms and downstream consequences of the aging processes remains a fundamental challenge, particularly in delineating primary drivers from compensatory responses and epiphenomena [1,2]. Current paradigms in aging research emphasize the fundamental roles of genomic integrity, transcriptional fidelity [3], and epigenetic regulation [4] as key determinants of lifespan and healthspan, providing potential targets for rejuvenation interventions [5]. However, a critical challenge in this field is distinguishing between organized, adaptive responses to aging and the accumulation of random, deleterious changes that contribute to functional decline [6,7].

This distinction becomes particularly relevant when considering the concept of biological entropy, the measure of disorder or randomness within biological systems [8]. While aging has long been conceptually associated with increased disorder, the systematic quantification of such disorder at the molecular level remains underexplored [9]. Information theory, originally developed by Shannon for communication systems [10], provides a mathematical framework to measure randomness and uncertainty that can be directly applied to biological data. In the context of gene expression, Shannon entropy quantifies the diversity and unpredictability of transcriptional patterns, offering a novel lens through which to examine age-related molecular changes [11].

Cancer and aging share a complex relationship [12]. On one hand, aging is the greatest risk factor for most cancers [13]; on the other hand, some scientists argue that aging actively suppresses carcinogenesis [14]. In fact, delving into the molecular layers the multidirectional relationship becomes more evident: some processes move in opposite directions, such as stemness [15,16] and cellular senescence signatures [17], while others, like the accumulation of mutations [18], inflammation [19], and loss of gene expression identity [20], progress in the same direction. Interestingly, there is a trend for mechanisms that go in the same direction to be more pronounced in cancer. This dynamic relationship highlights the need for precise approaches to measure aging and its role in cancer, and in this regard, entropy-based metrics alongside aging clocks can be extremely useful tools for disentangling adaptive from maladaptive molecular changes [21].

Transcriptomic and epigenetic clocks have revolutionized age-related research, providing more precise measurements and showing associations with aging acceleration/deceleration, age-related diseases, and mortality [22–24]. However, the complete understanding of the molecular mechanisms driving these changes in gene expression and epigenetic patterns during aging is still lacking. In this regard, recent studies explored how stochastic processes can partially explain the epigenetic clocks and thus changes in methylation patterns during aging [25–27]. The relationship between epigenetic and transcriptomic entropy is fundamentally interconnected through the regulatory hierarchy of gene expression. Since the primary function of the epigenome is to control transcriptional programs, stochastic changes in DNA methylation and histone modifications during aging should logically propagate to create corresponding disorder in gene expression patterns. This cascade effect suggests that transcriptomic entropy may serve as a downstream readout of epigenetic dysregulation, potentially capturing the cumulative impact of random epigenetic drift on cellular function. While epigenetic clocks measure the regulatory potential encoded in chromatin modifications, transcriptomic entropy directly quantifies the functional output—the actual gene expression patterns that determine cellular phenotypes and responses to environmental stimuli. Therefore, measuring transcriptomic entropy provides a complementary and potentially more functionally relevant metric of age-related molecular disorder that bridges the gap between epigenetic alterations and their phenotypic consequences. This approach may help elucidate the mechanistic links between the stochastic processes underlying epigenetic aging and their ultimate impact on cellular function. In this study, we leverage information theory to measure the randomness of the transcriptome in thousands of human and mouse samples to explore the role of expression entropy in cancer and aging.

## RESULTS

### Aging-related transcriptomic entropy variation is tissue-specific

First, we calculated Shannon entropy, a measure of randomness and uncertainty from information theory, for all GTEx and TCGA samples. As shown in Supplementary Figure 1A, the entropy distribution reveals an intriguing pattern: among the five tissues with the highest entropy (cervix uteri, uterus, nerve, testis, and ovary), four are sexual tissues; however, in contrast, none of the five tissues with the lowest entropy (pancreas, blood, kidney, heart, and liver) are. Additionally, we also observed a relatively large variation in entropy across different tissues, in comparison to primary tumor entropy (which we will discuss later) which is, on average, higher levels (Supplementary Figure 1B). Next, we correlated the median entropy of GTEx tissues with tissue turnover using data (only for human tissues) from Richardson et al. [28]. We observed a negative correlation (Supplementary Figure 1C), indicating that tissues with low entropy have a slower renewal rate. This suggests a potential link between entropy and cellular proliferation. This finding is consistent with the methylation entropy described by Vaidya et al. [29].

Afterwards, to investigate the relationship between entropy and transcriptional aging (see Methods), we performed a direct correlation between these two parameters (Figure 1A). We observed that the variation in entropy with aging is highly tissue-specific, with tissues such as brain, blood, and stomach showing a decline in entropy during aging, while others, like adipose tissue, salivary gland, and skin, exhibit an increase.

**Figure 1.**
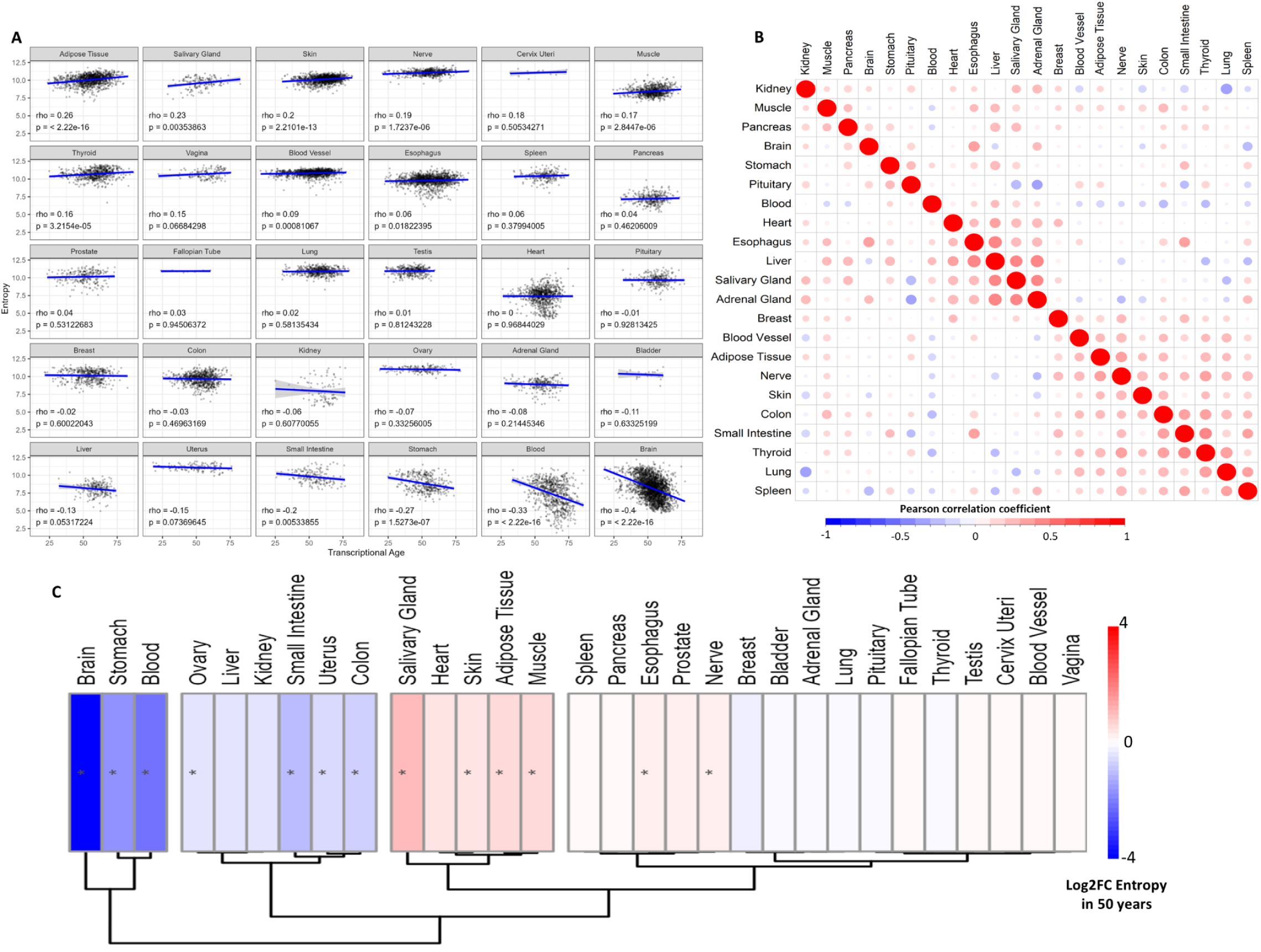
Transcriptomic entropy during human aging. A. Correlation trend between entropy and transcriptional age in human tissues. B. Correlation matrix of entropy values across different tissues from the same individual. C. Heatmap of entropy variation with transcriptional age, results from the adjusted linear model. * p-value < 0.05.

Exploring this tissue specificity, we build a correlation matrix (Figure 1B) to examine the association between entropy across different tissues from the same individual. We observed that, despite a general trend of positive correlations, there is a considerable number of negative and neutral correlations, indicating a lack of association between some tissues. Furthermore, there are only a few strong positive correlations with rho values above 0.4 (Liver x Esophagus, Salivary and Adrenal Glands, Salivary x Adrenal Gland, and Small Intestine x Thyroid). This supports the notion that the randomness of gene expression is a process that varies significantly across tissues, even within the same individual.

Finally, since it is well known that parameters like death and sex can influence gene expression (and consequently entropy) [30], we conducted a linear model analysis where we adjusted the variation in entropy with the variables available in GTEx (see Methods). In Figure 1C, we present a heatmap showing entropy variation over 50 years of transcriptional age. We can observe three main groups of tissues: those where entropy decreases (brain, stomach, blood, ovary, liver, kidney, small intestine, uterus, and colon), those where entropy increases (salivary gland, heart, skin, adipose tissue, muscle, and, to a lesser extent, esophagus and nerve), and those where entropy remains relatively unchanged (all other tissues).

We repeated the same analysis but now considering chronological age, and although the fold change scale was smaller, we observed essentially the same result (Supplementary Figure 2A). These findings surprisingly indicate that transcriptomic entropy does not uniformly increase with aging; rather, it is tissue-specific, showing a decrease or remaining stable in many tissues.

We also calculated the entropy for the brain, kidney, and liver for samples of dozens of mammals using data from Tyshkovskiy et al. [31] and Liu et al. [32] to check if the parameter is associated with the maximum lifespan of the species (Supplementary Figure 3). We found no significant association between transcriptomic entropy and maximum lifespan, at least in the species analyzed (although there seems to be a weak positive trend in the kidney). Interestingly, tissue entropy appears to be relatively at the same level among different species, which supports the idea of tissue specificity observed previously.

### Transcriptomic entropy is associated with age-related processes

Since we did not observe a unidirectional relationship between age and transcriptomic entropy, we sought to further investigate whether this stochastic process could play a role in aging. To do this, we first correlated entropy with phenotypes known to be associated with aging: stemness, cell proliferation (both of which tend to decrease with age), senescence, and the accumulation of somatic mutations (which tend to increase during aging).

Figure 2A-D shows the global correlations, where we observe strong positive correlations with proliferation and senescence (rho > 0.6) and a significant negative correlation with stemness (rho < -0.4). Furthermore, for approximately 7,000 samples with mutation data, we found a trend toward a positive correlation with entropy (rho > 0.25).

**Figure 2.**
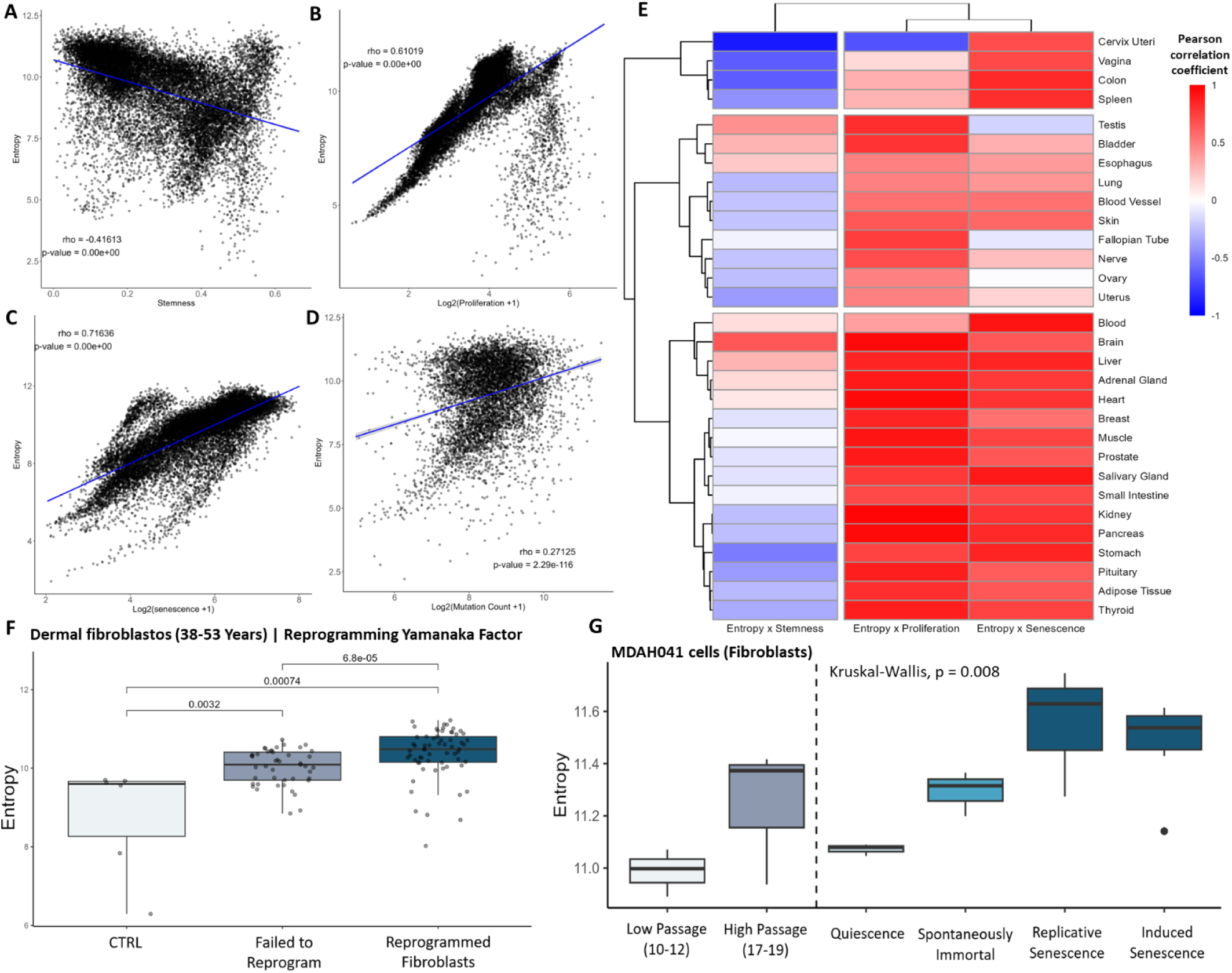
Transcriptomic entropy is associated with age-related processes. A-D. Correlation between entropy and stemness, proliferation, senescence and mutation count, respectively. **E.** Heatmap of tissue-specific Pearson correlation coefficient between entropy and stemness, proliferation and senescence. **F.** Comparison of entropy in fibroblasts treated with Yamanaka Factors: control, failed reprogramming, and successful reprogramming (data from Gill et al[36]). **G.** Impact of proliferation and senescence on entropy in MDAH041 fibroblasts (data from Purcell et al [37]). The p-value represents the global statistical significance.

In Figure 2E, we show the correlation coefficients in a tissue-specific heatmap for entropy versus stemness, proliferation, and senescence. We observe a more tissue-specific association with stemness, showing moderate values, which may indicate a more complex relationship, at least in adult tissues. For proliferation and senescence, we see a pattern of strong correlations across most tissues. Interestingly, all exceptions to this pattern (i.e., negative or neutral correlations) are reproductive tissues, which have different dynamics compared to somatic tissues [33]. Finally, for the samples with mutation data, we also generated a tissue-specific heatmap, where the pattern largely remained the same, highlighting a positive trend between somatic mutations and entropy in most tissues (Supplementary Figure 2B).

After that, in an attempt to understand possible mechanisms associated with transcriptome entropy, we studied similar parameters but now in cell lines. First, we examined changes to entropy during cellular reprogramming induced by Yamanaka factors, a promising rejuvenation strategy. As seen in Figure 2F, untreated fibroblasts have lower entropy than treated ones. Even more fascinating: fibroblasts that were treated but failed to reprogram (the authors of the original data assessed this process through cell membrane markers) exhibit less entropy than those successfully reprogrammed. Next, we analyzed the entropy of mouse fibroblasts reprogrammed by a chemical cocktail with 2 (2C) or 7 compounds (7C) (Supplementary Figure 2C). We can see that, although 2C has more entropy than 7C (which was more effective in reprogramming in the original paper [34]), chemical reprogramming also increases the entropy of the transcriptome compared to the control, corroborating the previous result. In other words, returning the cell to a more stem/embryo-like state seems to increase entropy. Along these lines, we used data from Cardoso-Moreira et al. [35] and observed that in most tissues analyzed, there is more entropy in the pre-birth period, which supports the previous finding (Supplementary Figure 4).

We also analyzed the variation of entropy with calorie restriction (CR) and rapamycin treatment in both mice and humans, and, at least in the tissues analyzed, we did not observe a difference compared to the control, which suggests that this variation may be specific to reprogramming and not to any anti-aging intervention (Supplementary Figure 2D-G).

Finally, we wondered about the strongly positive relationship between entropy, proliferation, and senescence: two intuitively opposite processes. To explore this, we analyzed fibroblast data where we can measure both the effect of proliferation and senescence. Despite the small sample size, which prevents pairwise comparisons, Figure 2G shows that fibroblasts with higher passage clearly exhibit more entropy than those with lower passage numbers. Additionally, quiescent cells (i.e., those that have exited the cell cycle) have less entropy than immortalized cells, and most notably, senescent cells (whether replicative or induced) show the highest entropy in the system. This corroborates previous findings and suggests that both proliferation and senescence directly affect transcriptome entropy.

This set of results suggests that: I- entropy changes along biological processes linked to aging, and II- age-related processes in opposite directions seem to positively influence transcriptome disorder. This partially explains the tissue specificity observed in the previous section: the different dynamics of tissues during aging likely determine the entropy patterns found.

### Increase in transcriptomic entropy during carcinogenesis

The results from the previous section suggest that an embryonic state and higher proliferation are associated with high transcriptomic entropy, which may resemble an oncogenic transformation. To verify whether the increase in entropy is associated with carcinogenesis, we first compared normal samples from GTEx with adjacent normal tissue and primary tumors from TCGA. We found that, in most cases, cancer exhibits higher entropy than normal tissue (Figure 3A). However, there are some important exceptions: lung tumors, where normal tissue has higher entropy, and thyroid (THCA) and uterine (UCEC) tumors, where the primary tumor has less entropy. Additionally, most results place adjacent normal tissue at an intermediate level of transcriptomic disorder, which aligns with the existing literature [38]. This suggests that, at least in general, the primary tumor has more entropy than normal tissue. One could argue that since GTEx and TCGA process their data slightly differently, this might impact the results. Considering this, we compared only the TCGA data, using adjacent normal tissue and primary tumors from the same patients (paired data), and obtained essentially the same results, reinforcing our conclusion (Supplementary Figure 5) of entropy-dependent mechanisms driving carcinogenesis.

**Figure 3.**
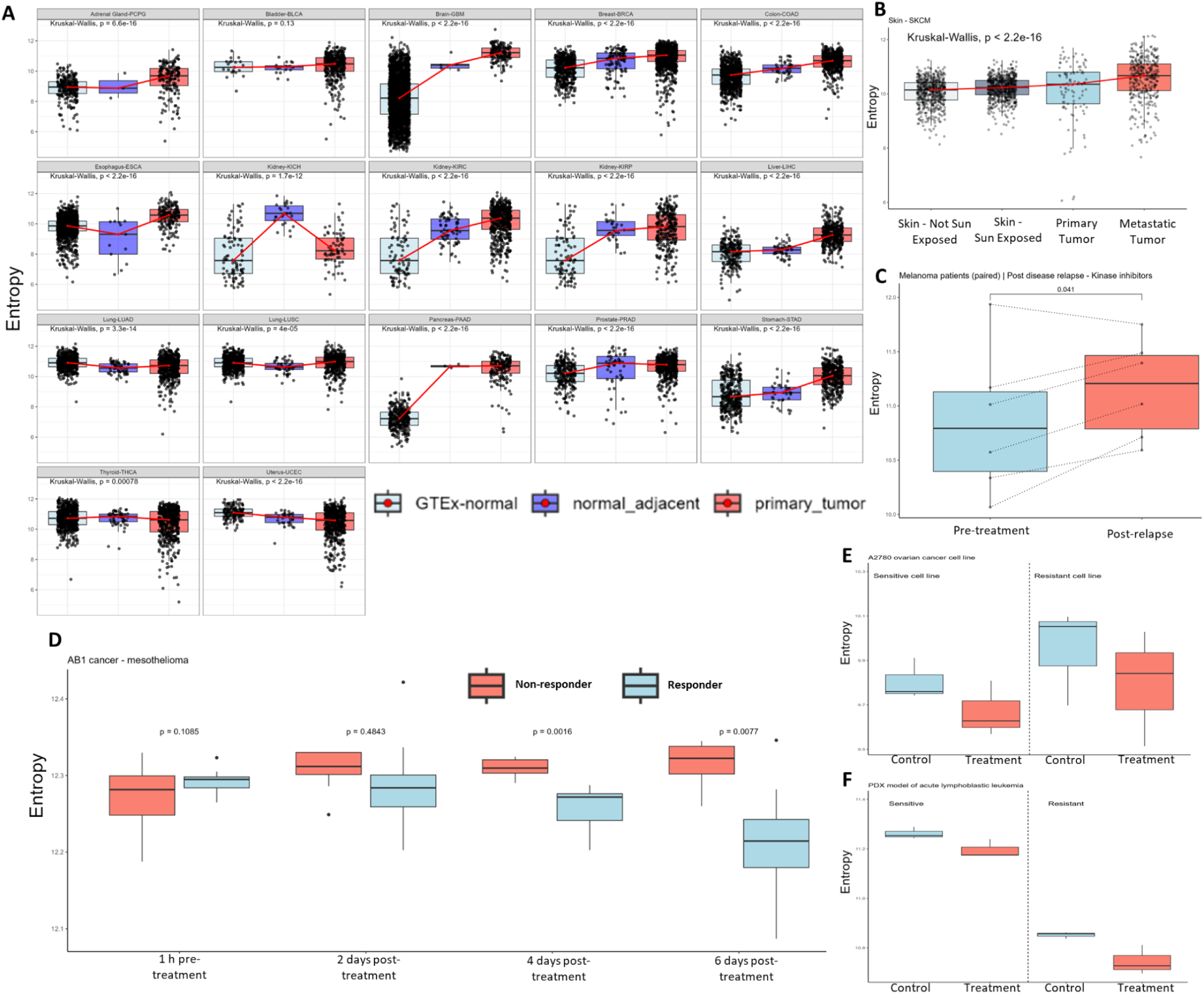
Entropy and carcinogenesis. **A.** Comparison of entropy between normal tissue (GTEx), adjacent tumor tissue, and primary tumor tissue (TCGA). **B.** Entropy in non-sun-exposed and sun-exposed normal skin (GTEx), and primary and metastatic melanoma (SKCM-TCGA). Red lines connect the medians to better visualization **C.** Comparison of entropy pre-treatment and post-acquired resistance (paired melanoma samples treated with RAF or RAF+MEK inhibitors) data from Wagle et al. [39] D. Comparison of entropy between responders and non-responders in mesothelioma (mice) treated with immune checkpoint blockade, data from Zemek et al. [40]. E. Ovarian cancer cell line treated with cisplatin, sensitive and resistant cells, data from Berman et al. [41]. F. Xenograft of ALL treated with dexamethasone and/or decitabine, sensitive and resistant samples, data from Jing et al[42].

Next, we wonder whether entropy plays a role in the progression of cancer to metastatic tumors. TCGA only provides an adequate number of metastatic samples for melanoma. We then took advantage of melanoma data (primary and metastatic) alongside normal skin, both sun-exposed and non-exposed (from GTEx), as sun exposure is the major carcinogenic factor for this disease. In Figure 3B, we observe an almost linear increase in entropy from non-exposed skin (healthy) to a late stage as metastatic tumors. This reinforces that stochastic transcriptomic processes are strongly associated with carcinogenesis.

Furthermore, when analyzing paired samples pre-treatment and post-acquired resistance, we see that 5 out of 6 patients had an increase in entropy post-relapse, with the only exception being the patient who already had the highest entropy in the population (both pre- and post-treatment) (Figure 3C). This suggests that the treatment-resistance dynamic may impact transcriptomic entropy.

Exploring this, we first checked whether differences in entropy are associated with treatment response. We used murine models of mesothelioma, where we found that over time, tumors that respond to treatment have significantly lower entropy than non-responders (Figure 3D). We observed a similar (albeit more modest) result in renal cell carcinoma (Supplementary Figure 6A). A direct interpretation is that the anti-tumor effect of treatment (when effective) reduces transcriptome disorder by eliminating tumor cells.

Another interpretation is that the treatment is selecting for resistant cells, and this effect is more pronounced when the treatment is effective; that is, by decreasing cellular heterogeneity, we would reduce overall entropy. With this in mind, we investigated the direct effects of treatments on cultured cells, where the environment is controlled and we could better observe what the treatment does to the tumor cell population. In Figure 3E, it can be observed that ovarian cancer cell lines treated with cisplatin exhibit a reduction in entropy post-treatment, whether they are resistant or sensitive.

Similarly, when we further delve into xenografts of acute lymphoblastic leukemia (ALL) derived from patients, it also shows decreased entropy after treatment, regardless of whether they are sensitive or resistant (Figure 3F). Interestingly, in ALL, the resistant cells have lower entropy, but it is worth noting that in this case, we are dealing with primary resistance rather than acquired. Finally, we used cellular models of androgen-dependent prostate cancer (LNCaP), where the cells were cultured with and without a testosterone analogue (DHT) (Supplementary Figure 6B). In this experiment, the authors treated the cells with different blockers of the androgenic pathway, and in all cases, the untreated controls (VEH) had the highest entropy in the system, suggesting that treatment with any drug, whether or not androgens are present, reduces entropy. This set of results reinforces the idea that treatment is a selective factor and, therefore, partially acts to decrease the randomness of gene expression signatures.

Finally, we examined the association of entropy with the number of somatic mutations and non-synonymous TMB. Similar to the results from GTEx, we observed a trend of positive associations in both cases, but with some exceptions (Supplementary Figure 7A-D). This suggests that transcriptome entropy is partially associated with genetic instability in both tumors and normal tissue.

### Entropy predicts cancer survival

To determine whether entropy is associated with tumor prognosis, we first examined the association with patient age, as aging alters the molecular landscape of tumors [43]. In Supplementary Figure 8A, we observe a prevalence of negative correlations between patient age and their transcriptional entropy. Furthermore, the six results we considered significant (|rho| > 0.15, p-value < 0.05) reinforces the notion that older patients have lower entropy (Supplementary Figure 8B). This aligns with the idea that tumors in younger patients are more aggressive, as they must deal with a host that has sharper tumor-suppressive mechanisms. Therefore, this suggests that entropy may be associated with aggressiveness.

Subsequently, we assessed whether transcriptomic entropy could predict survival. To do this, we first divided the primary tumor samples into two groups based on entropy (see Methods) and calculated a hazard ratio for overall survival (OS) and progression-free interval (PFI). In Figure 4A, we observe that, in most cases, higher entropy is associated with poorer prognosis. In Figure 4B, we show the six most significant results for OS, where five tumors (ACC, KICH, LGG, LIHC and MESO) exhibit lower survival in the group with higher transcriptomic entropy, with KIRC being the only exception. In Supplementary Figure 9, we present the other significant results (p < 0.05), confirming this trend, with seven out of ten tumors showing worse survival with higher entropy.

**Figure 4.**
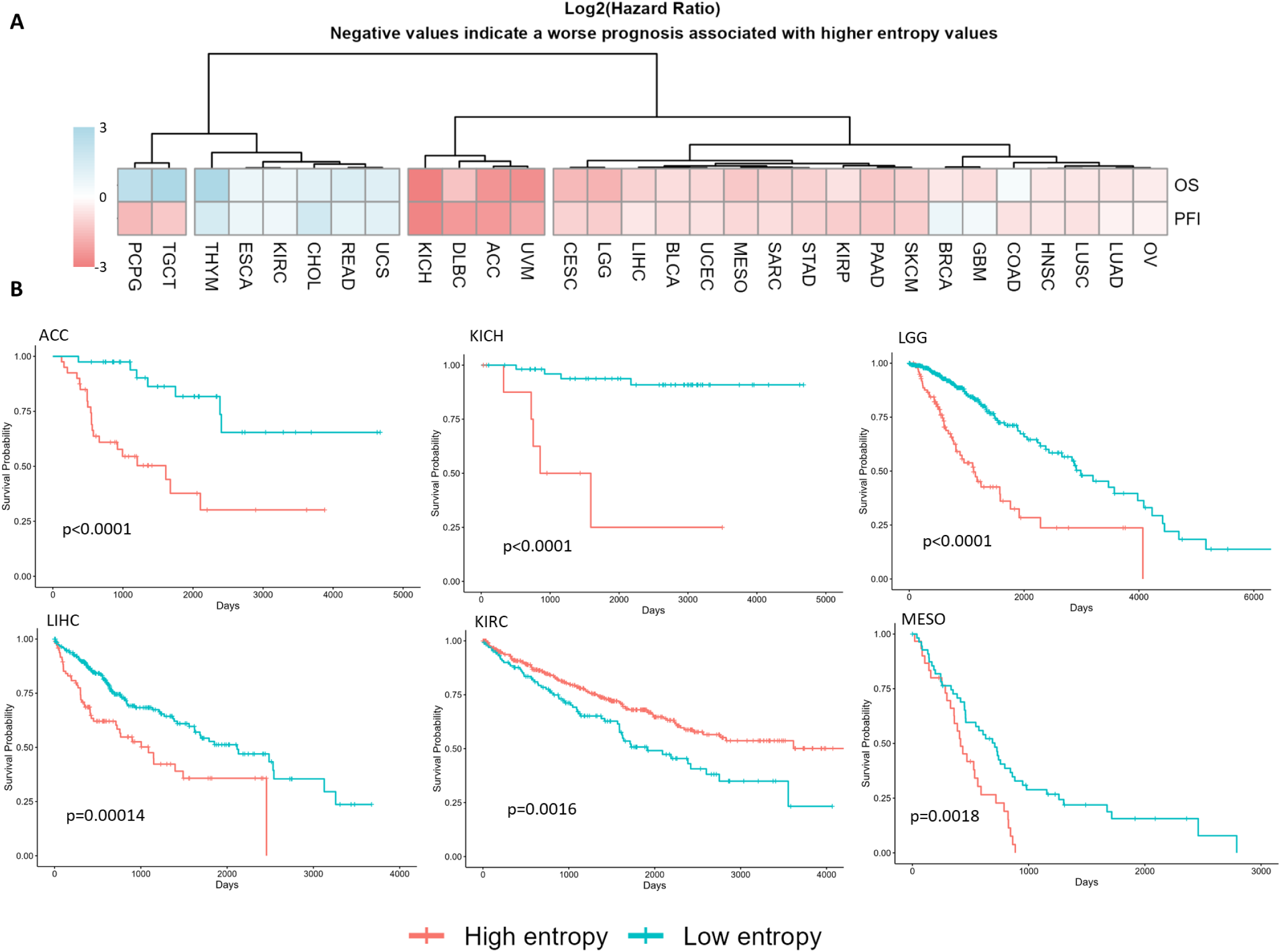
Entropy and cancer survival. **A.** Heatmap of hazard ratio for overall survival (OS) and progression-free interval (PFI).We exclude tumors without enough events in OS and/or PFI. **B.** Kaplan-Meier curves of the most significant results (lowest p-values) for overall survival from the previous heatmap. Data from primary tumors.

We repeated the Kaplan-Meier analysis for PFI, and in summary, the significant results showed that in 14 tumor types (ACC, CESC, DLBC, HNSC, KICH, KIRP, LGG, LIHC, LUSC, PAAD, PCPG, SARC, TGCT, and UVM), high entropy is associated with poorer prognosis, whereas in 5 other cases (CHOL, KIRC, THCA, THYM, and UCS), it appears to be protective (data not shown). This reinforces the idea that, in most cases, higher entropy in gene expression is associated with increased tumor aggressiveness.

### Enrichment analysis links entropy with cancer processes

Our previous results suggest that intrinsic factors (such as proliferation, senescence and oncogenic transformation) and extrinsic factors (such as sun exposure and cancer treatment) could modulate transcriptome entropy. However, is there a gene signature associated with cells exhibiting higher entropy? To gain insights into this, we performed a differential expression analysis to identify protein-coding genes that show significant variation in expression relative to entropy changes in both normal tissues (GTEx) and primary tumors (TCGA).

When comparing the overlap of up- and downregulated genes between GTEx and TCGA, we found that of the 94 genes present in at least two lists, only two showed opposite directions (i.e., upregulated in GTEx and downregulated in TCGA, and vice versa). This suggests that despite differences in samples, contexts, and processing, there is a conserved expression pattern in more stochastic samples (Figure 5A).

**Figure 5.**
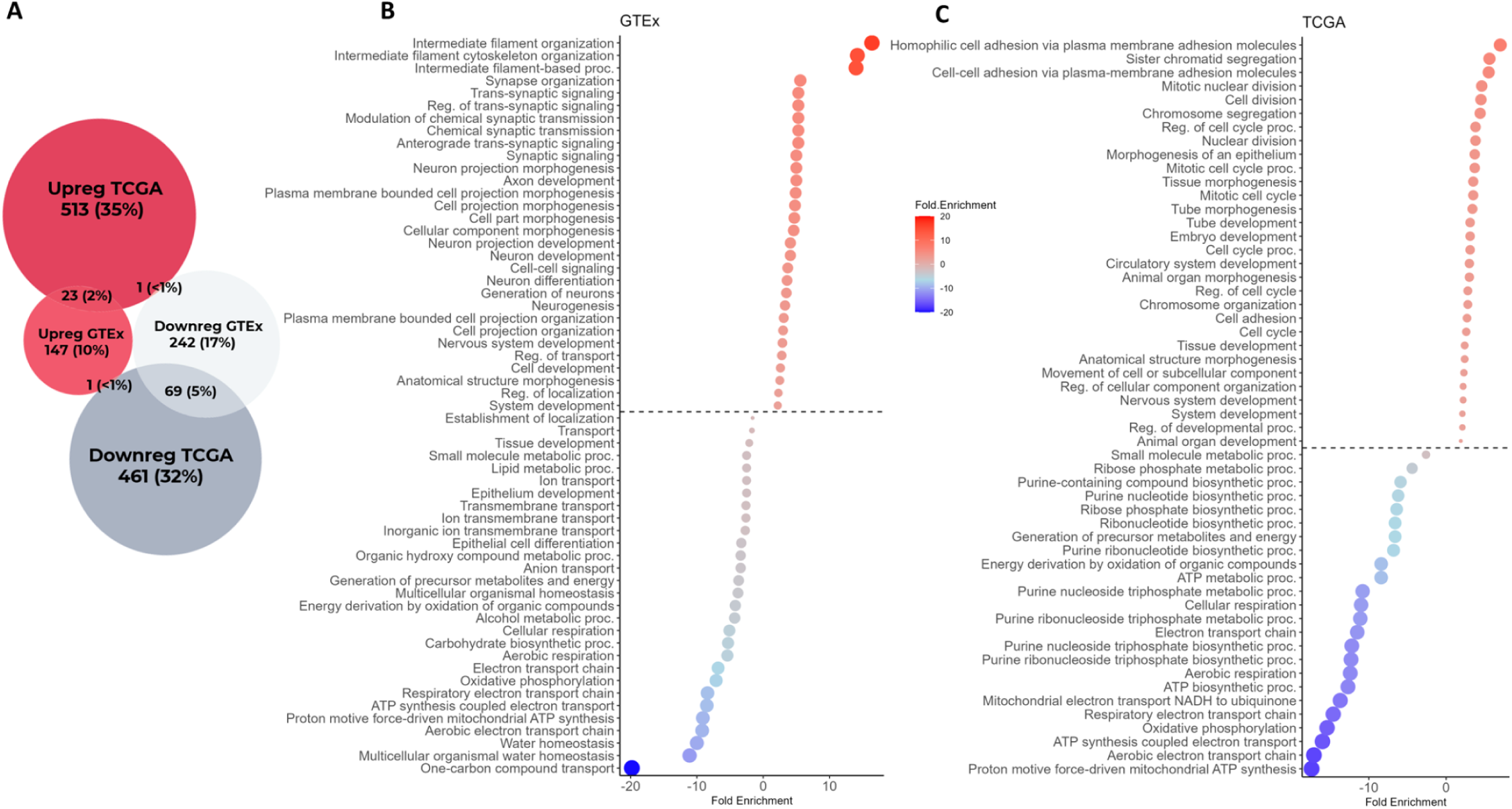
Enrichment analysis for protein-coding genes with expression variation associated with entropy. A. Euler diagram of differentially expressed genes in relation to entropy in GTEx and TCGA samples B-C. Enrichment of biological processes for upregulated (red) and downregulated (blue) genes from GTEx and TCGA, respectively.

To reinforce this notion, we conducted a functional enrichment analysis for biological processes (Gene Ontology) and observed an interesting result: despite the differences, both normal and tumor tissues show enrichment for developmental and morphogenesis-related genes, and depletion for genes involved in metabolism and cellular respiration (Figure 5B-C).

This leads us to important considerations: I – although we are quantifying randomness, there appears to be an expression signature associated with entropy variation. II – Particularly for GTEx samples, the enriched processes seem indicative of dedifferentiation, as we observe developmental processes and shifts in cellular metabolism in adult somatic tissues (in a Warburg effect-like manner) [44]. This again reinforces the notion that increased entropy is associated with aging and associated-carcinogenesis.

Next, as we previously observed an increase in entropy with cellular reprogramming (Figure 2F), we repeated the same approach to investigate the biological processes associated with reprogramming, and whether they correlate with those enriched in the GTEx and TCGA entropy analyses. In Supplementary Figure 10, we found that the processes of upregulated genes are mainly associated with the cell cycle, similar to those of the TCGA, which makes sense since cellular reprogramming can lead to cancer. On the other hand, the depleted processes, although there is some overlap in transport-related functions with GTEx, are quite different. This may suggest that during reprogramming, despite the increase in entropy, the activated/deactivated genes differ from those in normal tissue, with an obvious overlap (increased proliferation) with cancer.

### Other age-related diseases also exhibit a trend toward an increase in transcriptomic entropy

Since aging is a major risk factor not only for cancer but also for other chronic diseases, we hypothesized that the increase in transcriptomic entropy would be involved in their progression too. GTEx classifies its tissue samples as normal; however, most samples undergo pathological analysis, and many include pathologist reports noting alterations (e.g., inflammation, hyperplasia). Thus, although these samples are considered healthy, they often exhibit pathological changes. We compared the median entropy between affected and unaffected samples across available tissues and found that most tissues show increased entropy in affected samples (Figure 6A), suggesting that elevated stochastic processes at the transcriptional level may be associated with the development of pathological conditions. Next, we compared mouse models of age-related diseases to assess whether there are alterations in transcriptomic entropy in tissues affected by these diseases. In Figure 6B, all four datasets used demonstrate higher entropy in diseased tissues compared to healthy controls.

**Figure 6.**
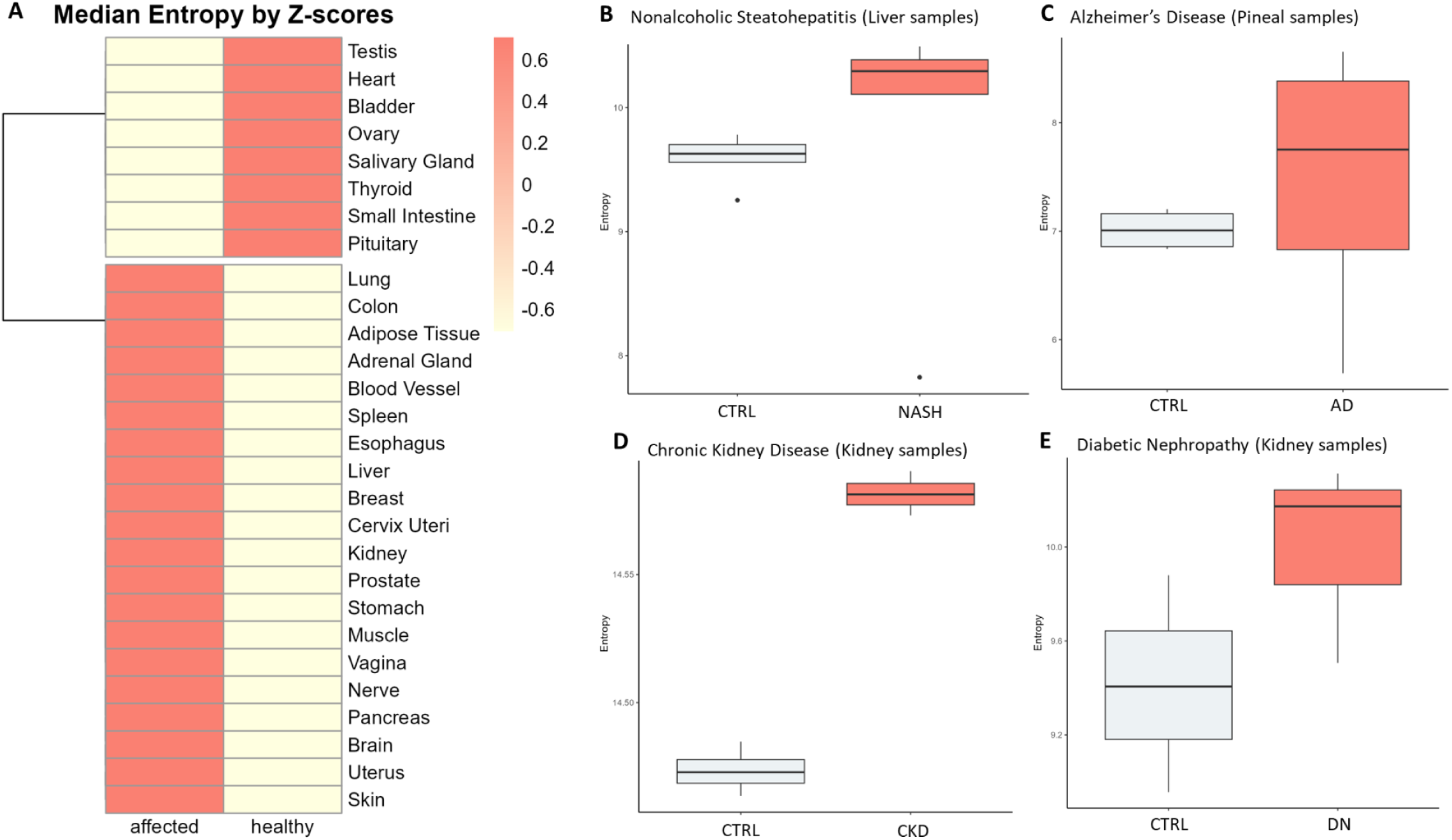
Transcriptomic entropy may be related to age-related diseases. A. Z-score heatmap comparing the median entropy between affected and healthy samples. B. Boxplot showing differences in entropy between control and age-related diseases. CTRL = control

These last analyses, while not comprehensive due to the limited number of samples, suggest that increased entropy may also be associated with other age-related diseases beyond cancer, opening up interesting opportunities for future studies.

## DISCUSSION

The concept of increased disorder impacting aging, longevity and associated diseases is not new and is intuitively accepted due to its logical consistency and some supporting evidence [6,7,45]. For instance, as previously mentioned, some recent studies clearly demonstrate that there is a random component in the variation of methylation during aging [25–27]. A recent study even suggests that methylation entropy is a biomarker of aging [9]. Here, it is important to emphasize that epigenetic alterations primarily result in changes in gene expression, which are associated with aging [46]. Furthermore, the concept of transcriptional drift and noise suggests that variations in expression can be both random and non-random [47]. However, a study on the subject failed to detect a robust increase in transcriptional noise in aged tissues [48], indicating a lack of consensus on the matter and the need for further exploration. Here, we comprehensively studied the application of Shannon entropy to thousands of mammalian samples in an effort to understand the role of transcriptomic entropy (i.e., disorder and randomness) in aging and associated diseases, particularly cancer.

First, we analyzed how entropy changes during aging in human tissues, using transcriptional and chronological ages. Surprisingly, most tissues maintain stable levels of entropy or even reduce it. Moreover, the levels of entropy (and their variation with aging) appear to be tissue-specific. Consistently, the predominance of entropy reduction during aging, and tissue specificity has been observed in scRNA-seq studies involving various mouse tissues [49]. When we delved into certain age-associated processes, we found that opposing processes (senescence and proliferation) have a positive effect on entropy, which is similar to the conclusion made by Okada in a recent study [50], which may partially explain the tissue-specificity of entropy.

Another surprising finding was that transcriptome entropy, an intuitively detrimental phenomenon, increases during cellular reprogramming, an anti-aging therapy. This also reinforces the mutual relationship between cancer and aging: we know that reprogramming can lead to cancer [51], and all our results indicate that the increase in entropy is associated with the progression of most analyzed tumors, which is aligned with the literature [21]. We also observed that anti-cancer treatment impacts entropy levels, which was expected since it is well-established that therapies affect gene expression profiles [52–54] and, consequently, our measure of transcriptomic entropy.

Furthermore, our cancer results suggest that transcriptome entropy may serve as a prognostic biomarker (associated with survival) and could aid in understanding (and potentially predicting) treatment responses, paving the way for its application in personalized medicine, as previously proposed by Conforte et al [11]. We also observed an increase in entropy in other age-related diseases, which could be further explored in future studies, as stochastic molecular processes have already been associated with these types of conditions [55,56].

Although Shannon entropy is directly associated with randomness, we were able to find a similar expression pattern in the samples from GTEx (normal) and TCGA (primary tumor). More interestingly, these patterns indicate a cellular dedifferentiation and a metabolic alteration, hallmarks of carcinogenesis [57]. The accumulation of senescent cells, despite being non-dividing cells, can stimulate carcinogenesis through the senescence-associated secretory phenotype (SASP)[58,59], and our results suggest that senescence itself also increases transcriptomic entropy. Additionally, processes associated with dedifferentiation, which we observe in the upregulated genes in relation to entropy, are directly linked to the phenotypic plasticity necessary for tumor progression [16,60,61]. This leads us to wonder whether entropy could serve as a marker of carcinogenesis and, importantly, if it can be used to detect cancer in its early stages, as well as being a potential biological link between aging and carcinogenesis.

Entropy is originally a concept from thermodynamics that measures the degree of disorder and randomness in a physical system. In this study, we applied Shannon entropy to the transcriptome, considering that we are measuring disorder and randomness; however, we cannot exclude the possibility that we are (even partially) measuring an increase in system complexity or heterogeneity (rather than disorder per se). In cancer, it is easy to understand that a tumor cell has more disordered and complex expression patterns simultaneously. However, in aging, these two parameters may counterbalance each other. For instance, it is reasonable to argue that a cell undergoing reprogramming is not "disordered," but rather there is an increase in expression complexity/heterogeneity that, through a direct entropy measure, may appear as disorder. For example, a study demonstrated that activation of oncogene-induced senescence pathways in cells is accompanied by a progressive, time-dependent increase in gene expression heterogeneity [62]. This finding aligns with our observation of entropy increase during senescence and suggests that, in certain contexts, our entropy measure captures the diversity of gene expression within the cellular population. This is significant because we can consider: I - there may exist a beneficial range of entropy as well as a deleterious one, which can vary from context to context. II - It is crucial that future studies delve deeper into the differentiation between randomness and biological complexity to better understand the biological mechanisms associated with aging processes.

It is important to discuss some limitations of this study. First, all our analyses were conducted using bulk RNA-seq, which, although easier to interpret and apply, may obscure whether changes in transcriptomic entropy occur due to alterations in the expression patterns of the same cell types or changes in cell proportions (thus "distorting" the initial gene expression). Conclusions drawn by Yang et al. [63] indicate that long-lived, high-turnover cells are more susceptible to age-related variations in transcriptomics, suggesting the first scenario. However, further studies, particularly utilizing scRNA-seq, need to delve deeper into this topic, as both scenarios may occur (or counterbalance each other) simultaneously, depending on the cell/tissue type. Additionally, the analyses conducted here have overlooked the retrotranscriptome, which comprises approximately 50% of our genome and has important implications for both cancer and aging [64]. In future work, we aim to understand whether the expression of retrotransposons influences transcriptomic entropy and what its impact is on cancer and aging. Finally, in a subsequent study, we aim to identify which genes contribute most significantly to the increase in entropy and to investigate the underlying reasons for this phenomenon. For instance, a recent study demonstrates that longer genes are more prone to accumulate random damage and alter their expression patterns [65], which is expected to be reflected in transcriptomic entropy. This is particularly relevant in the context of diseases, as it may help establish biomarker signatures.

In conclusion, our study demonstrates the tissue-specific nature of transcriptomic entropy during aging, revealing that different tissues exhibit distinct patterns of entropy variation. The association of entropy with key biological processes such as proliferation, senescence and cellular reprogramming provides new insights into the complexities of age-related processes. Importantly, our findings suggest that transcriptomic entropy could serve as a valuable prognostic biomarker in cancer, highlighting its potential role in understanding tumor progression and treatment responses. Additionally, the increased entropy observed in other age-related diseases, along with the biological processes associated with this rise, points to a broader relevance of this concept, which reinforces the need for further investigations into the connections between transcriptomic entropy, aging, and disease mechanisms.

## METHODS

### Data acquisition

For aging analyses in human tissues, RNA-Seq-based gene expression data from healthy samples were downloaded from the GTEx portal (https://gtexportal.org) in TPM and read counts (version 8) [66].

For human cancer analyses we utilized TCGA RNA-Seq data (from primary tumor, metastatic tumor and normal adjacent tissue) also in TPM and read counts, which we downloaded from GDC portal (https://portal.gdc.cancer.gov/). For both TCGA and GTEx, the raw data were aligned to the human genome GRCh38/hg38 by their respective consortia.

Samples without complete information (age, sex, survival,etc) were filtered out .

To support our findings, we also downloaded processed data from the GEO (https://www.ncbi.nlm.nih.gov/geo/) database, using expression data from humans and mice. If there was any genetic editing or treatment that could interfere with the analysis, only the control was used for the analyses.

The datasets used were: GSE60340 (senescence and proliferation), GSE165176 (reprogramming), GSE77940 (melanoma samples pre- and post relapse) GSE173201 (ovarian cancer cell line), GSE109947 (acute lymphoblastic leukemia xenograft), and GSE211781 (prostate cancer cell line), GSE153941 (AB1 and Renca mouse tumors), GSE103532 (kidney - CR), GSE230002 (muscle - CR). For mice data, we used RNA-seq data from: GSE246954 (chemical reprogramming), GSE120977 (NASH), GSE129586 (AD), GSE189377 (CKD), and, GSE197699 (DN) and GSE230002 (rapamycin treatment).

Finally, we downloaded human RNA-Seq data from pre- and postnatal samples by Cardoso-Moreira et al in FPKM [35].

### Shannon entropy

Shannon entropy is a measure of uncertainty or randomness in a system, widely used in various fields to quantify diversity. In the context of transcriptomics, it allows us to assess the diversity of gene expression across samples or genes [10]. By considering the distribution of gene expression levels, Shannon entropy provides a measure of how uniformly genes are transcribed within a given sample. A higher entropy value reflects greater diversity, while lower entropy suggests more uniform gene expression.

To calculate the transcriptome diversity for each sample the expression levels were normalized by dividing each value by the total sum of expression for that sample, resulting in the relative proportions of gene transcription. Using these proportions, we applied the Shannon entropy formula, where each proportion is multiplied by the base-2 logarithm of that same proportion. The final entropy value is the negative sum of these products, providing a measure of gene expression diversity within each sample. This calculation was performed individually for each sample, yielding the overall transcriptome randomness. The formula used is as follows:

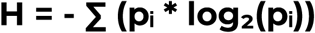

Where pᵢ represents the proportion of expression for the i-th gene in relation to the total gene expression in the sample. The summation (∑) is over all genes i in the sample, calculating the contribution of each gene to the overall entropy.

### Chronological and biological age

Since GTEx open-data only lists age ranges (i.e., 20–29, 30–39, 40–49, 50–59, 60–69 and 70-79), age was approximated to 25, 35, 45, 55, 65 and 75, respectively, as before [20]. This age was what we considered as “chronological age”.

To estimate biological age, we calculated the transcriptional age for each GTEx sample using the RNAAgeCalc tool [67] with the “predict_age” function, applying the following parameters: exprtype = ’counts’, signature = ’all’, and idtype = ’ENSEMBL’. This age was what we considered as “transcriptional age”.

### Linear model

To measure the entropy variation within aging (both chronological and transcriptional/biological) we applied a linear model similar to what was done previously [15].

For each tissue, fold change (FC) with age was calculated using the model below. To improve visualization, we multiplied the FC value by 50, resulting in the variation of entropy over 50 years. If any variable is not present in the tissue (e.g., sex for vagina or prostate), it is disregarded in the analysis. All information on the subjects was taken directly from the GTEx portal.

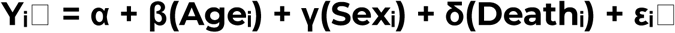

The variables are defined as follows:

- Yᵢ□: The entropy j in sample i.
- Ageᵢ: The age of sample i – continuous variable.
- Sexᵢ: The sex of sample i – categorical variable.
- Deathᵢ: The death classification of sample i based on the 4-point Hardy scale – categorical variable [68].
- εᵢ□: The error term for entropy i in sample j.

The variable age was used independently with both chronological and transcriptional age (see above). Linear model was generated using the R package limma, using the lmFit() function [69,70].

### Correlogram

To assess the entropy behavior between tissues across the same individual, we constructed a correlation matrix using Pearson coefficients (correlogram). For the few cases in which a specific subject had two or more samples from the same tissue, we used the average entropy to calculate the correlation matrix.

We build the plot using the R package ‘corrplot’ with hierarchical clustering option (https://github.com/taiyun/corrplot). In this analysis, we excluded sexual tissues and tissues with limited sample sizes, specifically: ’Testis’, ’Prostate’, ’Ovary’, ’Uterus’, ’Vagina’, ’Fallopian Tube’, ’Cervix Uteri’, and ’Bladder’, as these would interfere with the hierarchical classification.

### Correlation with mammalian maximum lifespan

To correlate transcriptome entropy with maximum lifespan, we used RNA-seq data from Tyshkovskiy et al. [31] and Liu et al. [32] , and longevity data from Human Ageing Genomic Resource [71]. We considered the median entropy for each species/tissue. The maximum lifespan was converted to log10 for better visualization. Outlier species, named in the plot, in each tissue were determined automatically using ggrepel (https://github.com/slowkow/ggrepel), with default parameters.

The species used in these analyses were: Acomys cahirinus, Arctonyx collaris, Atelerix albiventris, Balaena mysticetus, Balaenoptera acutorostrata, Balaenoptera borealis, Bos taurus, Callithrix jacchus, Callosciurus erythraeus, Canis lupus, Capra hircus, Castor canadensis, Cavia porcellus, Cryptomys damarensis, Cynopterus sphinx, Dasypus novemcinctus, Daubentonia madagascariensis, Delphinus delphis, Equus caballus, Erinaceus amurensis, Erinaceus europaeus, Felis catus, Heterocephalus glaber, Homo sapiens, Hystrix brachyura, Lemur catta, Macaca fascicularis, Macaca mulatta, Meles meles, Meriones unguiculatus, Mesocricetus auratus, Microcebus murinus, Monodelphis domestica, Muntiacus muntjak, Mus musculus, Mustela putorius, Mustela sibirica, Myocastor coypus, Myotis brandtii, Myrmecophaga tridactyla, Neovison vison, Nyctereutes procyonoides, Nycticebus coucang, Ornithorhynchus anatinus, Oryctolagus cuniculus, Pachyuromys duprasi, Paguma larvata, Pan paniscus, Pan troglodytes, Papio hamadryas, Peromyscus leucopus, Petaurus breviceps, Plecotus auritus, Procavia capensis, Pteromys volans, Rattus norvegicus, Rhinolophus ferrumequinum, Sarcophilus harrisii, Sciurus vulgaris, Suncus murinus, Sus scrofa, Tadarida brasiliensis, Tamias sibiricus, Trachypithecus francoisi, Tupaia belangeri, Ursus americanus, Vulpes lagopus, and Vulpes vulpes

### Correlation with stemness, proliferation, cellular senescence and mutation count

In the GTEx samples, we correlated entropy with biological parameters that are known to be associated with aging. For stemness, we used values previously calculated in previous study [15].

For proliferation, we employed the gene list provided in the study by Ramaker et al. [72]. In summary, the authors established and validated a proliferation index for healthy tissues and cells using a gene list from Venet et al. [73]. This gene list for constructing the proliferation index was identified from the top 1% of genes most positively correlated with the proliferation marker PCNA across 36 normal tissues. We applied this gene list and calculated the average expression of all genes within the proliferation list, obtaining a proliferation value per sample.

For senescence, we used the same strategy as proliferation but utilized the SenMayo gene list [74] which is one of the most widely used by the community for senescence-related analysis.

Finally, for mutation count, we used data from García-Nieto et al., who estimated somatic mutations for approximately 7,000 GTEx samples [75].

### Clinical and molecular features for cancer analysis

To associate tumor entropy with clinical parameters (survival and age), we used the curated data from Liu et al. for the TCGA samples [76]. TMB and mutation count data were downloaded from the cBioPortal, using the TCGA Pan-Cancer Study [77].

To construct Kaplan-Meier curves and heatmaps of hazard ratios, we utilized the survival [78] and survminer (https://github.com/kassambara/survminer) packages in R. We filtered data for each cancer type and identified optimal cutpoints for entropy using the surv_cutpoint function (based on log-rank test). Subsequently, we created survival objects and fitted Kaplan-Meier models to visualize overall survival (OS) and progression-free interval (PFI) probabilities. Finally, Cox proportional hazards models were applied to calculate hazard ratios based on the defined entropy groups.

### Differential expression analysis and functional enrichment

To identify differentially expressed genes in relation to entropy, we applied a linear model similar to the one described above for GTEx samples and primary tumors from TCGA. The main differences from before included filtering to evaluate only protein-coding genes and excluding the Death variable for TCGA, while considering chronological age as the Age variable in both cases.

Furthermore, to achieve pan-tissue/cancer results rather than tissue-specific outcomes, we adjusted the model for tissue (GTEx) or tumor type (TCGA).

Considering that the FC result reflects the variation in expression relative per unit of entropy, we defined differentially expressed genes as those with |Log2 FC| > 0.25 and FDR < 0.05. We used these four gene lists (Upregulated GTEx or TCGA, Downregulated GTEx or TCGA) and performed functional enrichment analysis using ShinyGO 0.8, with parameters set to species = human, FDR cutoff = 0.05, pathways to show = 30, and pathway database = GO Biological Process. To improve visualization, the Fold Enrichment of downregulated genes was multiplied by -1.

### Pathology slides

To classify GTEx samples as healthy or affected, we downloaded the pathology slide descriptions from the consortium (https://gtexportal.org/home/histologyPage). We manually classified all available samples as healthy if the Pathology Category was empty or labeled as "clean_specimens," "spermatogenesis" (for testis), "no_abnormalities," "tma," or "clean_specimens, no_abnormalities”. The other samples were classified as affected and represent alterations noted by the pathologist, such as fibrosis, congestion, atrophy, hyperplasia, inflammation, etc.

### Data Analysis

All data and statistical analyses in this study were conducted using base R functions and ggpubr (https://github.com/kassambara/ggpubr) . The graphs were built using the ggplot2 [79] and pheatmap (https://github.com/raivokolde/pheatmap) packages, unless otherwise specified.

## Acknowledgments

GAS was supported by a fellowship from Fundação de Amparo à Pesquisa do Estado de São Paulo (FAPESP) [2023/11499-8]. PAFG was supported by grants from Serrapilheira Foundation and FAPESP [2018/15579-8].

