## Supplementary Figure for "Transcriptomic entropy reveals tissue-specific patterns in aging and predicts cancer progression"

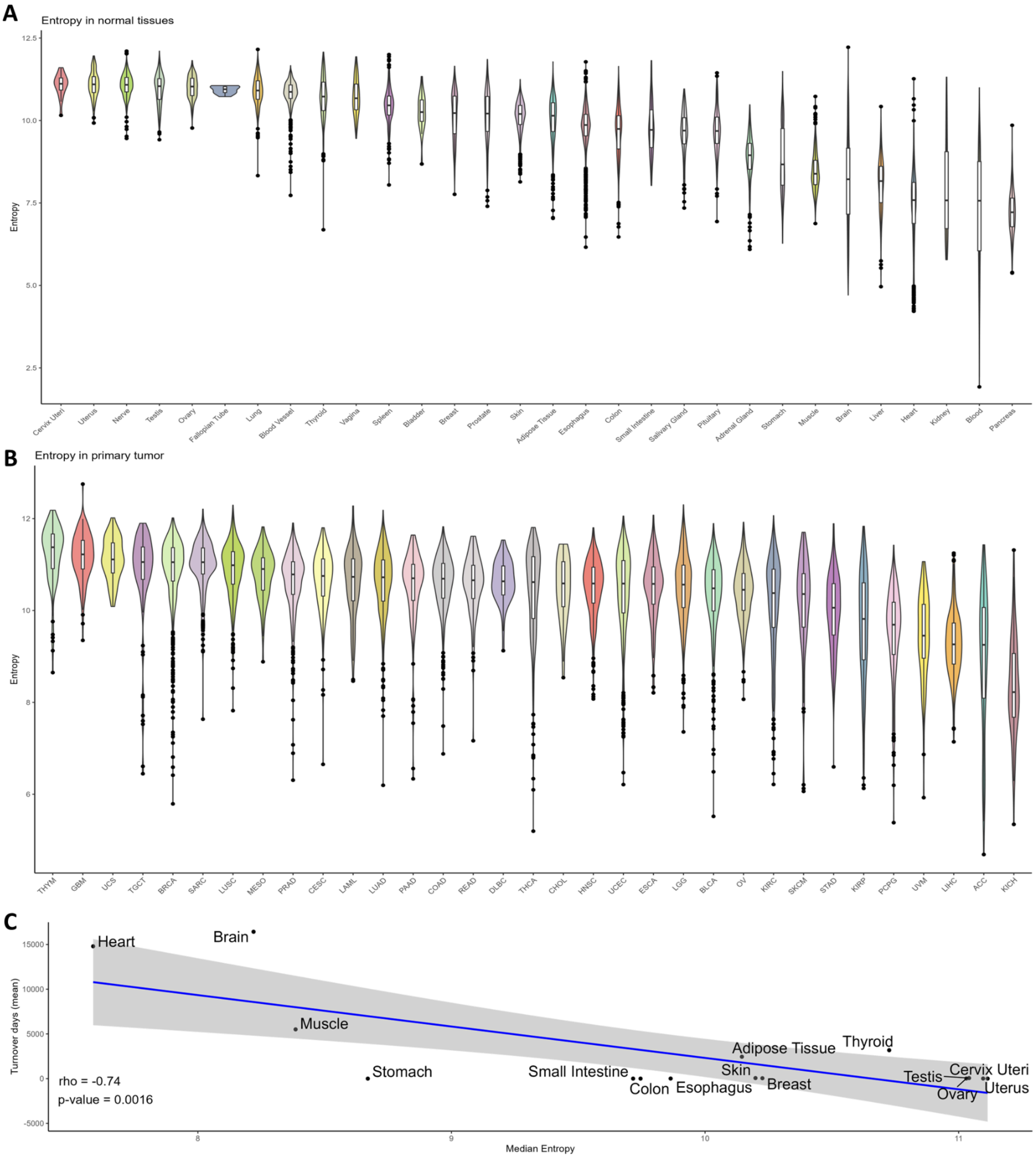


**Supplementary Figure 1. A-B.** Entropy distribution in GTEx and TCGA samples, respectively.
**C.** Correlation between tissue turnover and median entropy in normal (GTEx) tissues.

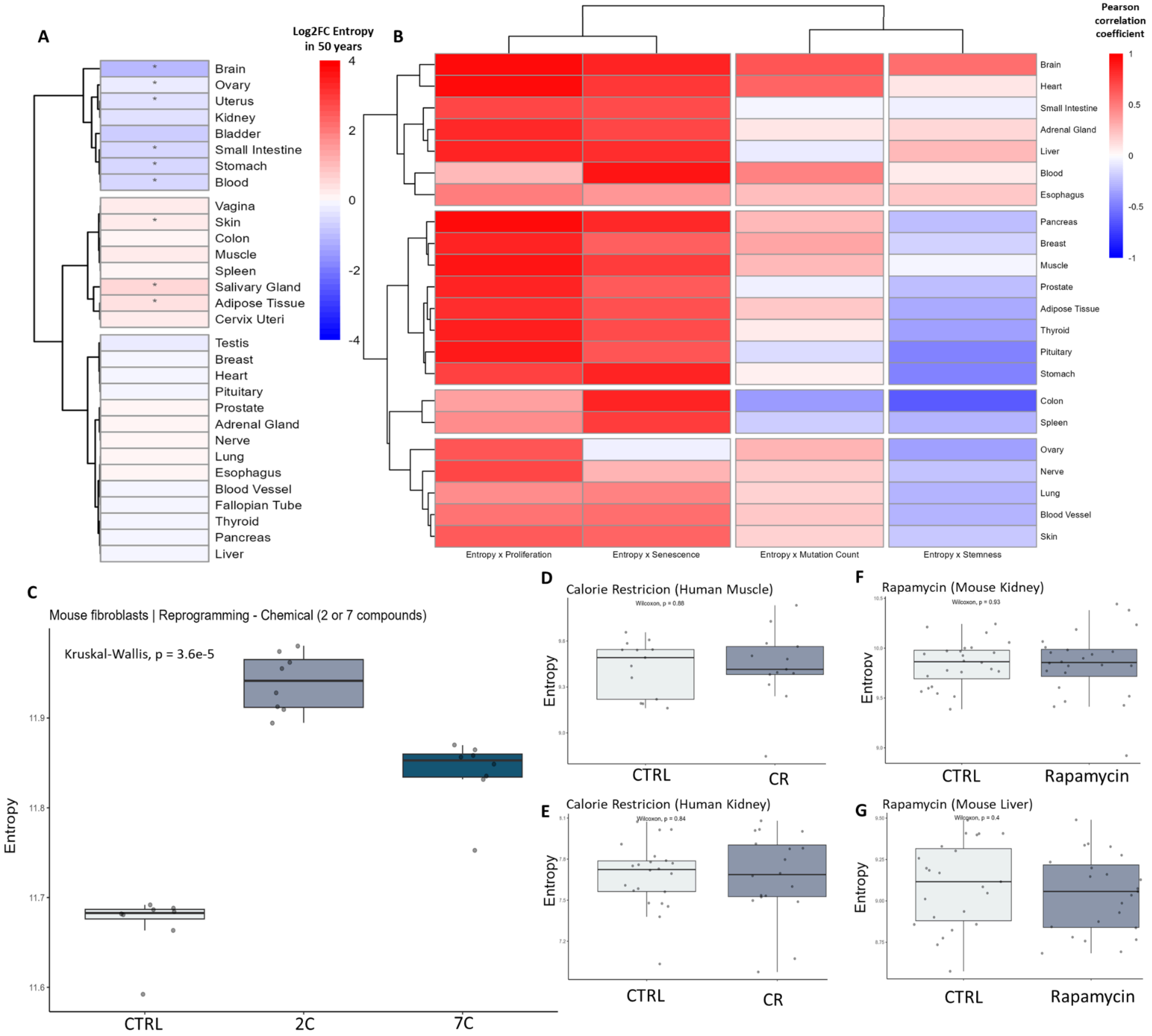


**Supplementary Figure 2. A** Heatmap of entropy variation with chronological age, results from the adjusted linear model. * p-value < 0.05. **B.** Heatmap of tissue-specific Pearson correlation coefficient between entropy and proliferation, senescence, mutation count and stemness for GTEx samples with mutation data. **C.** Entropy in chemical reprogramming from mouse fibroblasts using 7 (repsox, trans-2-phenylcyclopropylamine, DZNep, TTNPB, CHIR99021, forskolin, and valproic acid) or 2 compounds (repsox and trans-2-phenylcyclopropylamine). **D-E.** Entropy after calorie restriction (CR) in normal human tissues (kidney and muscle). **F-G.** Entropy after Rapamycin treatment in normal mouse tissues (kidney and liver). CTRL = Control

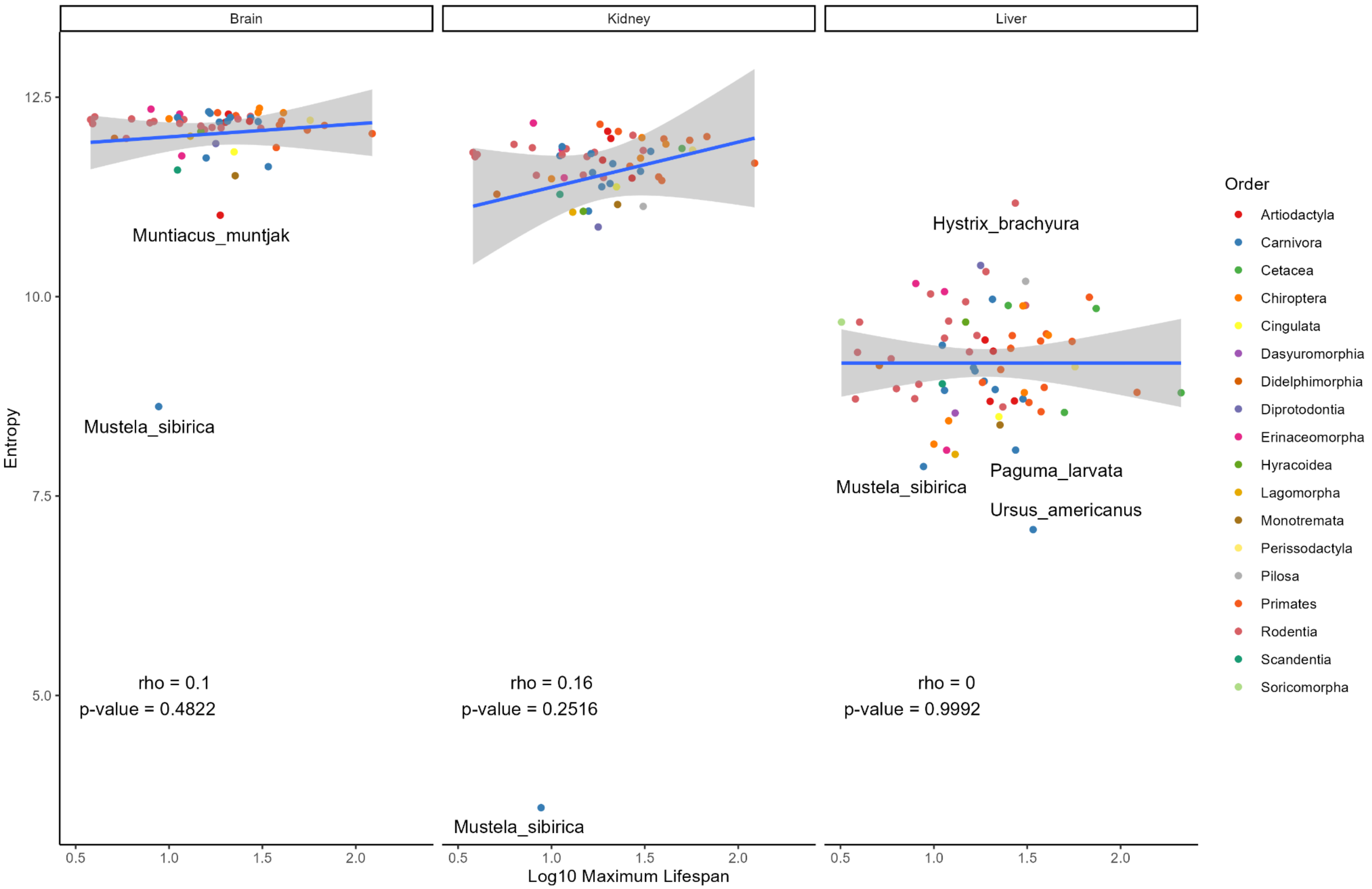


**Supplementary Figure 3.** Correlation between transcriptomic entropy and maximum lifespan in several mammals. Species that deviate from the pattern were named.

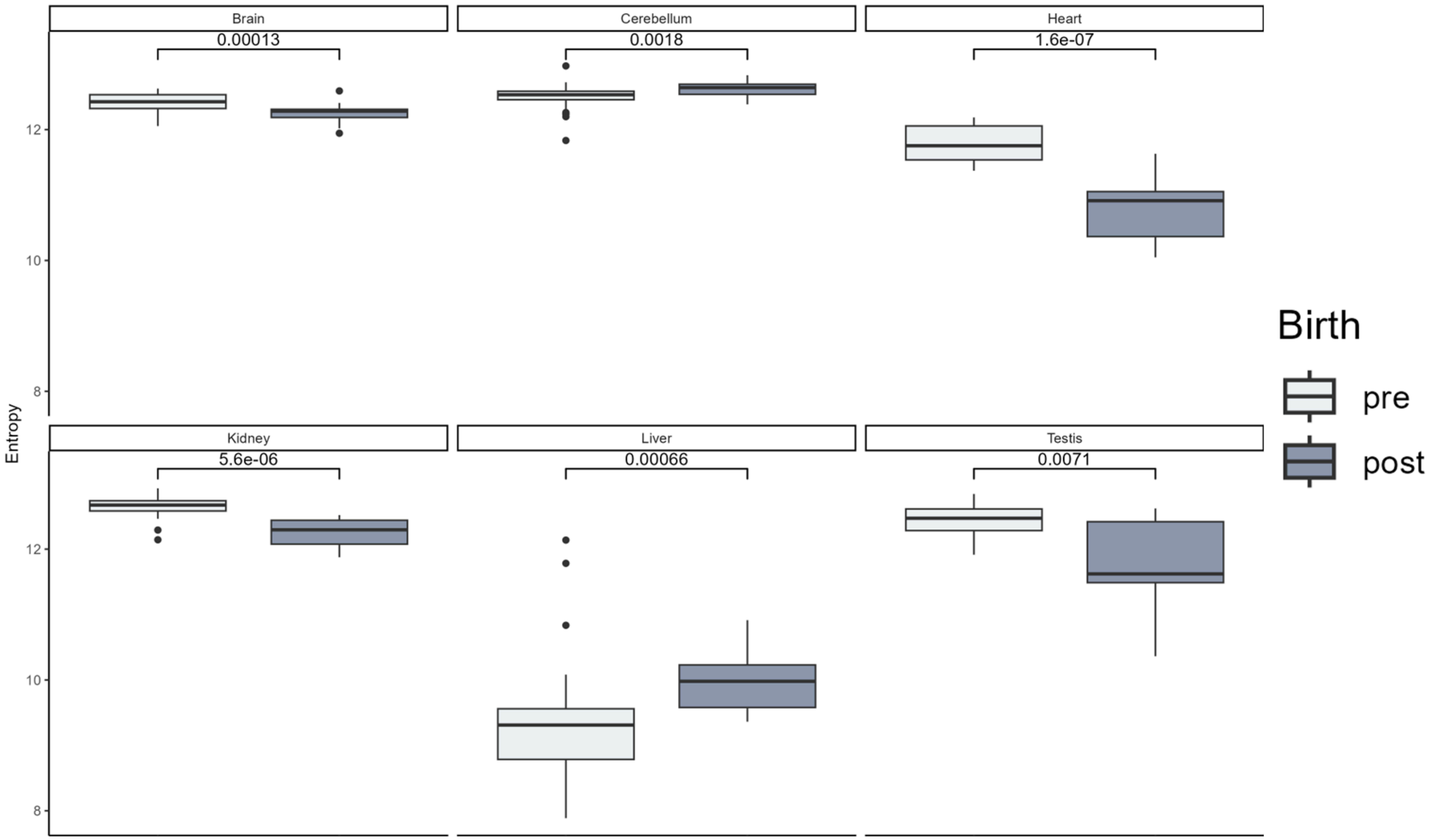


**Supplementary Figure 4.** Transcriptome entropy pre- and post- birth in human tissues.

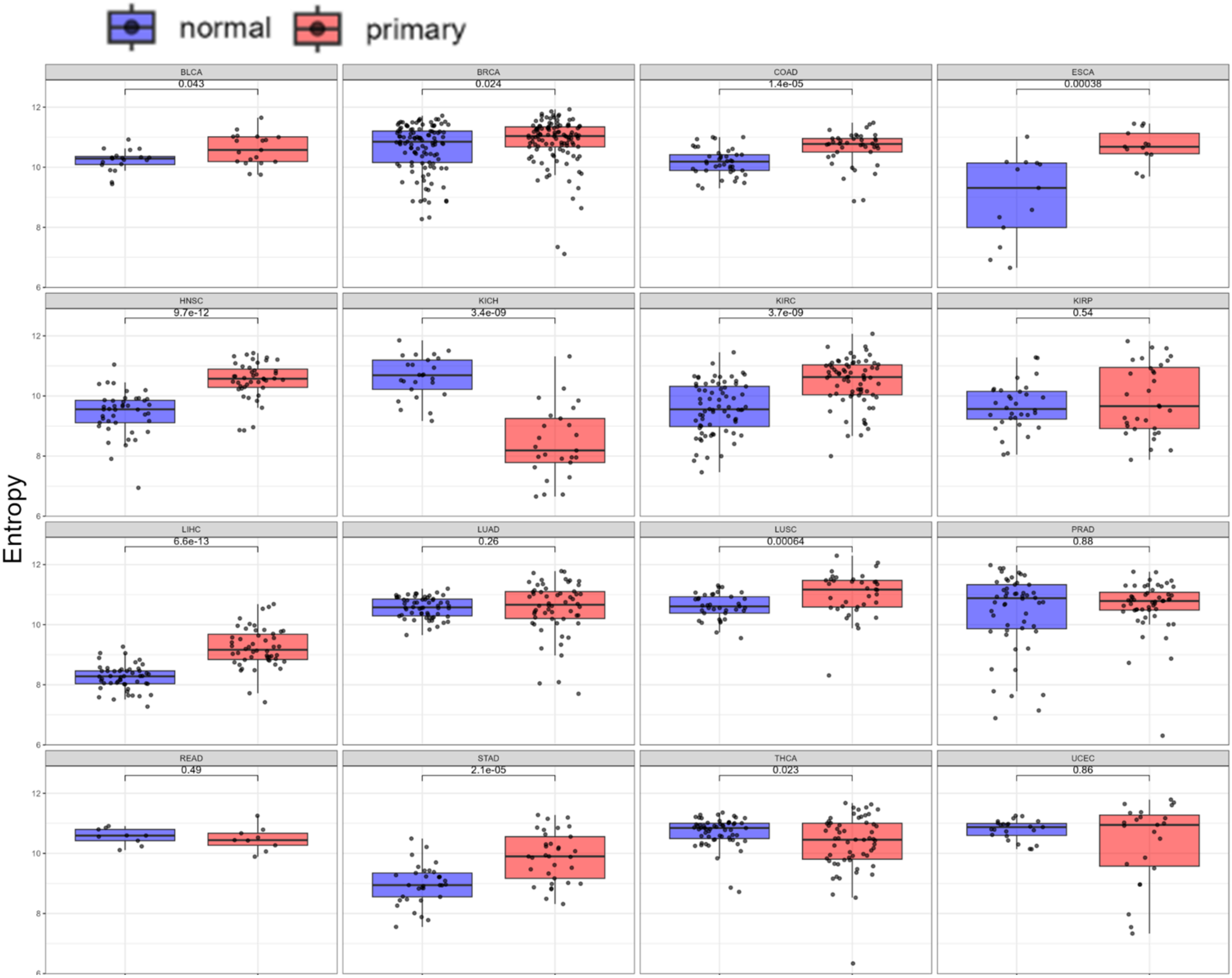


**Supplementary Figure 5.** Transcriptome entropy in normal (normal adjacent tumor) and primary (cancer) TCGA samples. Paired samples.

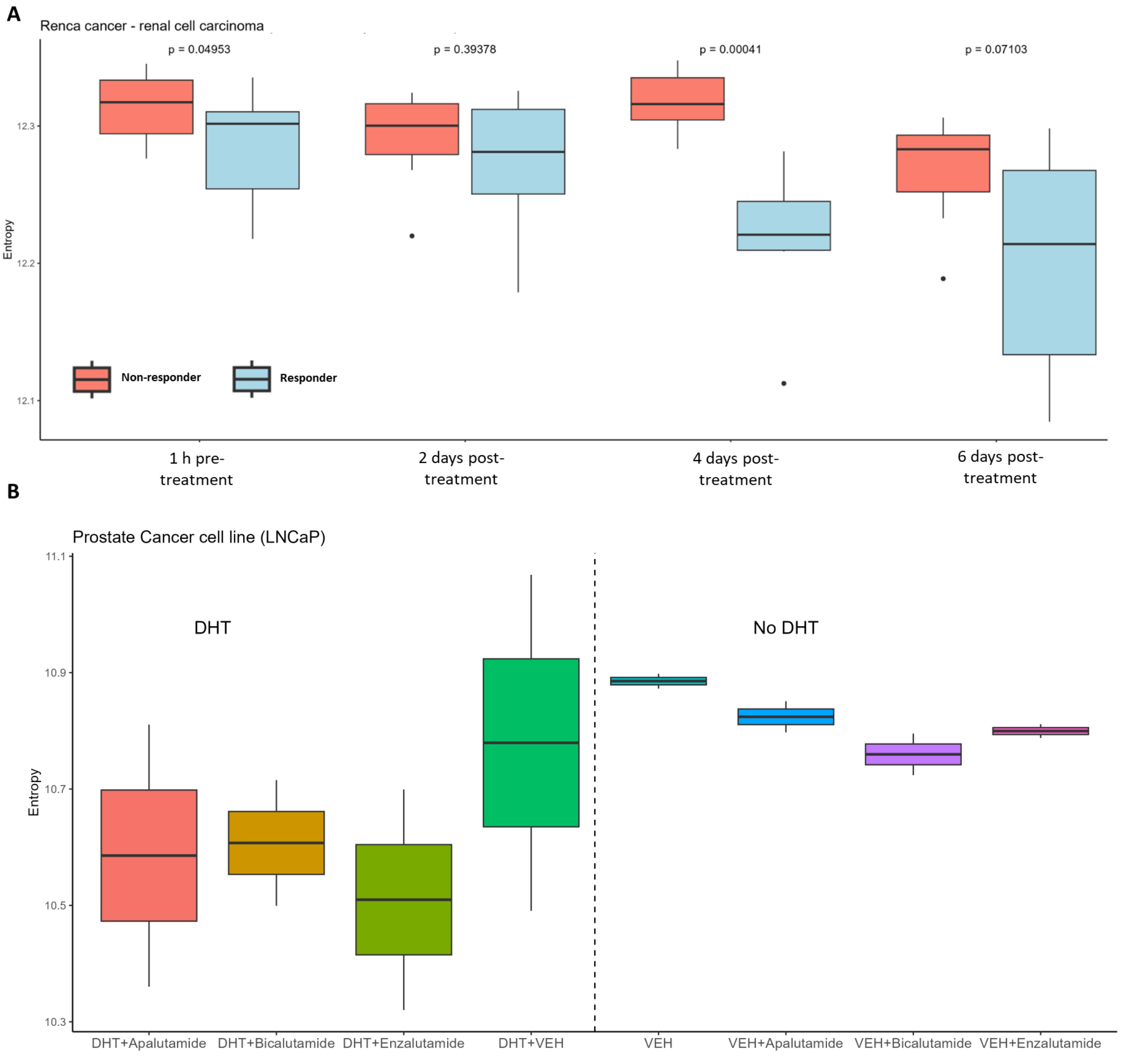


**Supplementary Figure 6. A.** Comparison of entropy between responders and non-responders in renal cell carcinoma (mice) treated with immune checkpoint blockade. **B.** Androgen-dependent prostate cancer cell line treated with different drugs that inhibit the pathway. DHT = Dihydrotestosterone. VEH = Vehicle used for control.

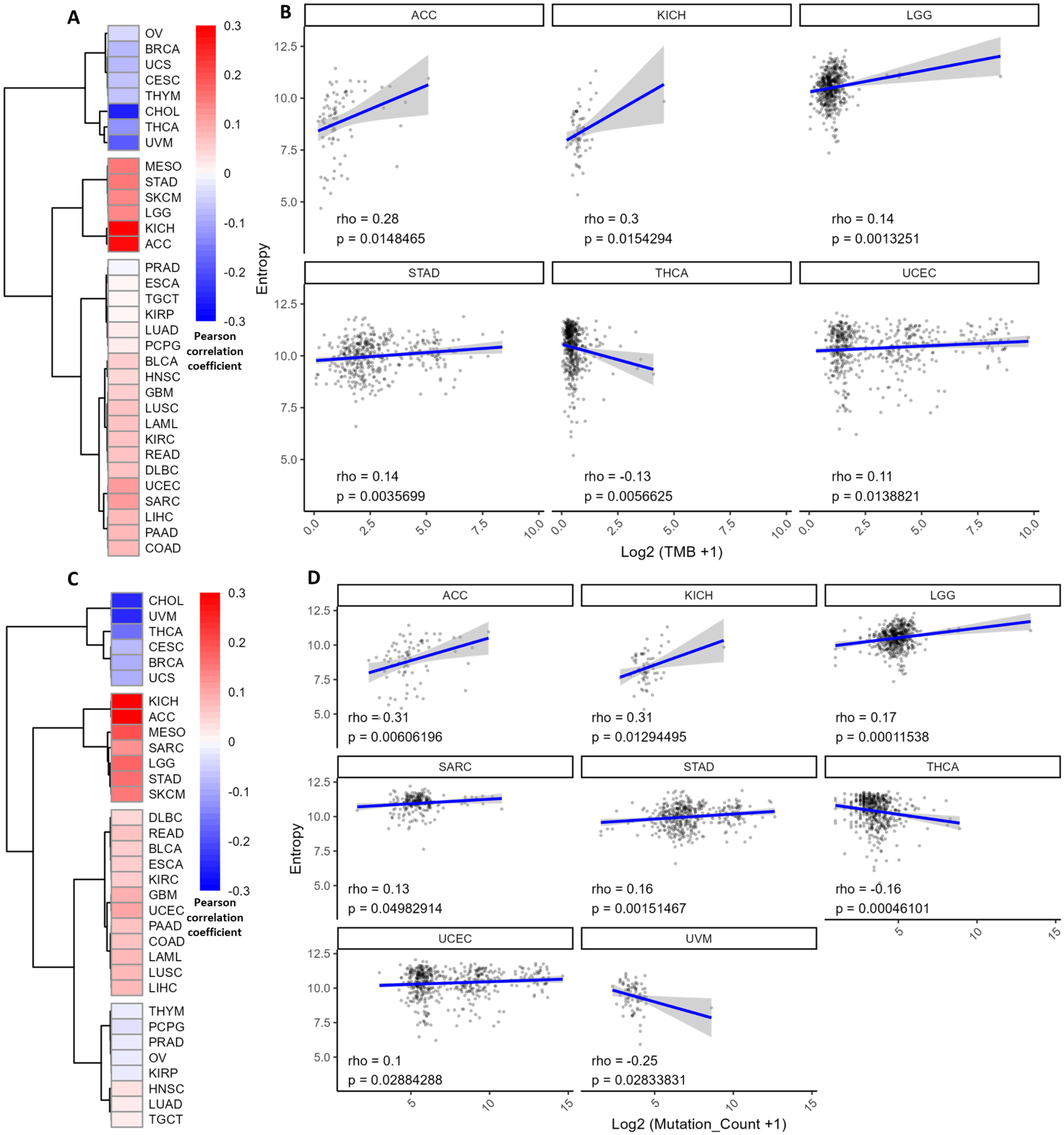


**Supplementary Figure 7. A.** Heatmap of Pearson' correlation between entropy and patient TMB (non-synonymous). **B.** Significant results from the previous heatmap. **C.** Heatmap of Pearson' correlation between entropy and all somatic mutations. **D.** Significant results from the previous heatmap. Data from primary tumors

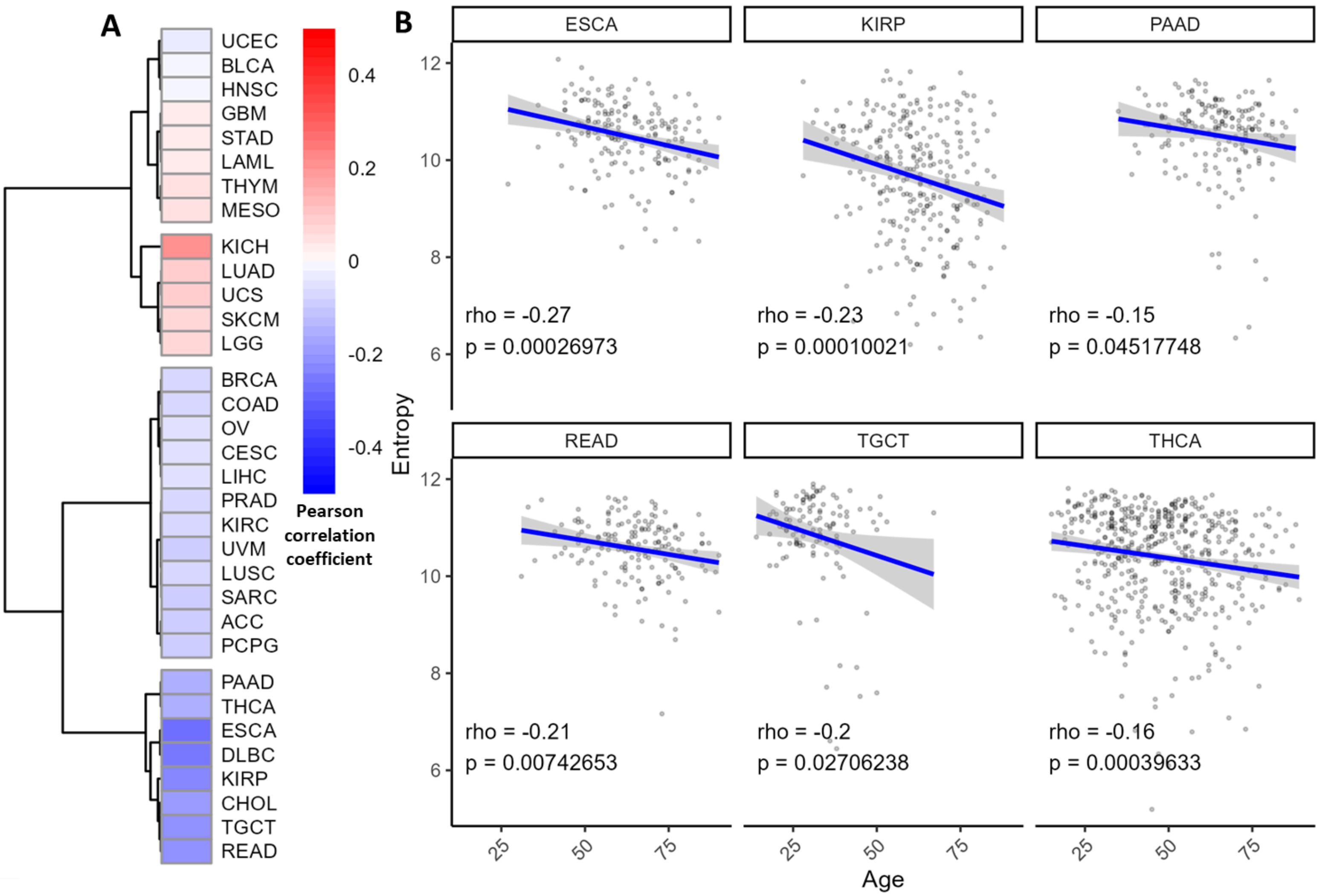


**Supplementary Figure 8. A.** Heatmap of Pearson' correlation between entropy and patient age. **B.** Significant results from the heatmap. Data from primary tumors

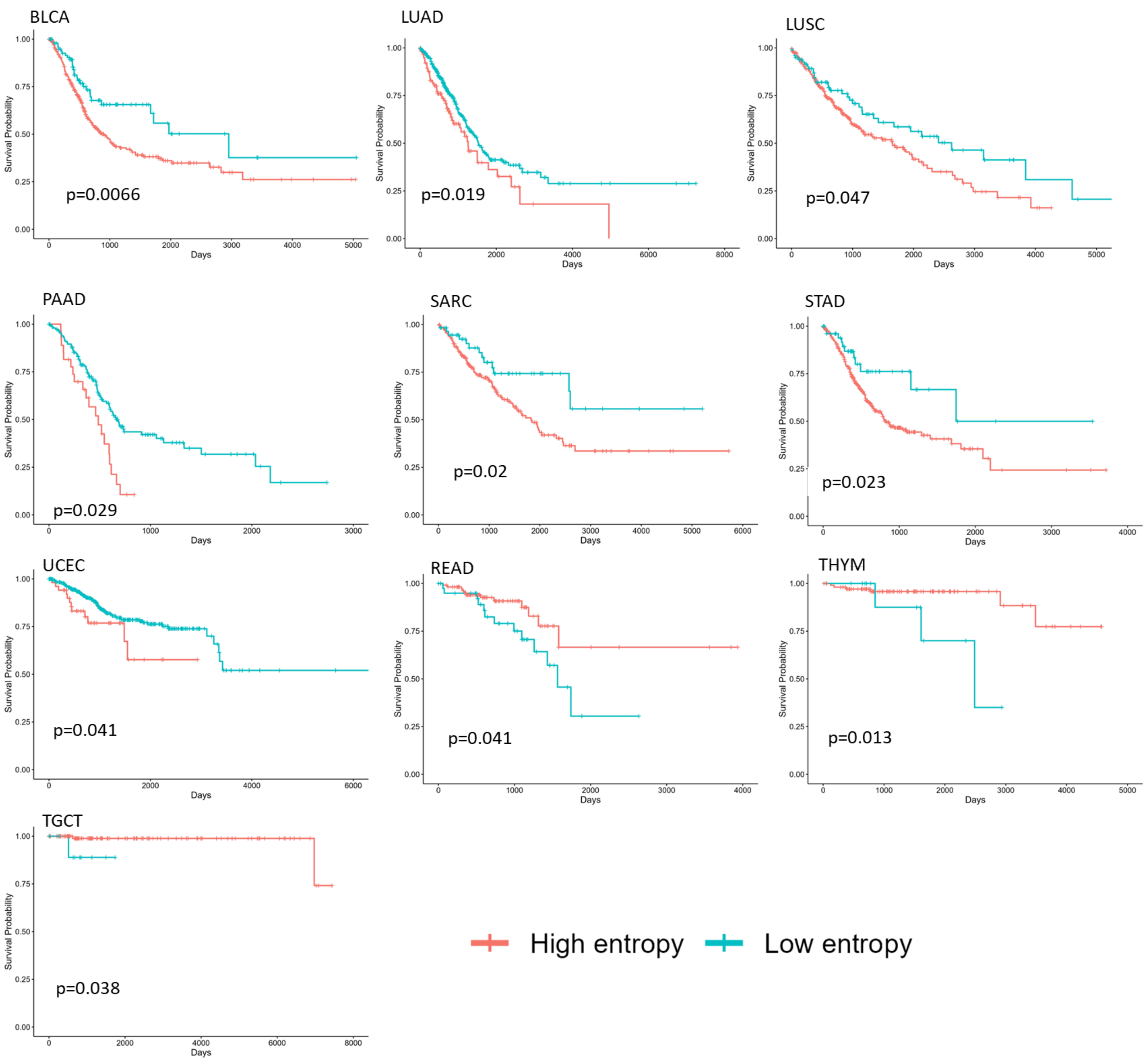


**Supplementary Figure 9.** Kaplan-Meier curves for overall survival for significant results ($p<0.05$) excluding those already shown in figure 4B. Data from primary tumors

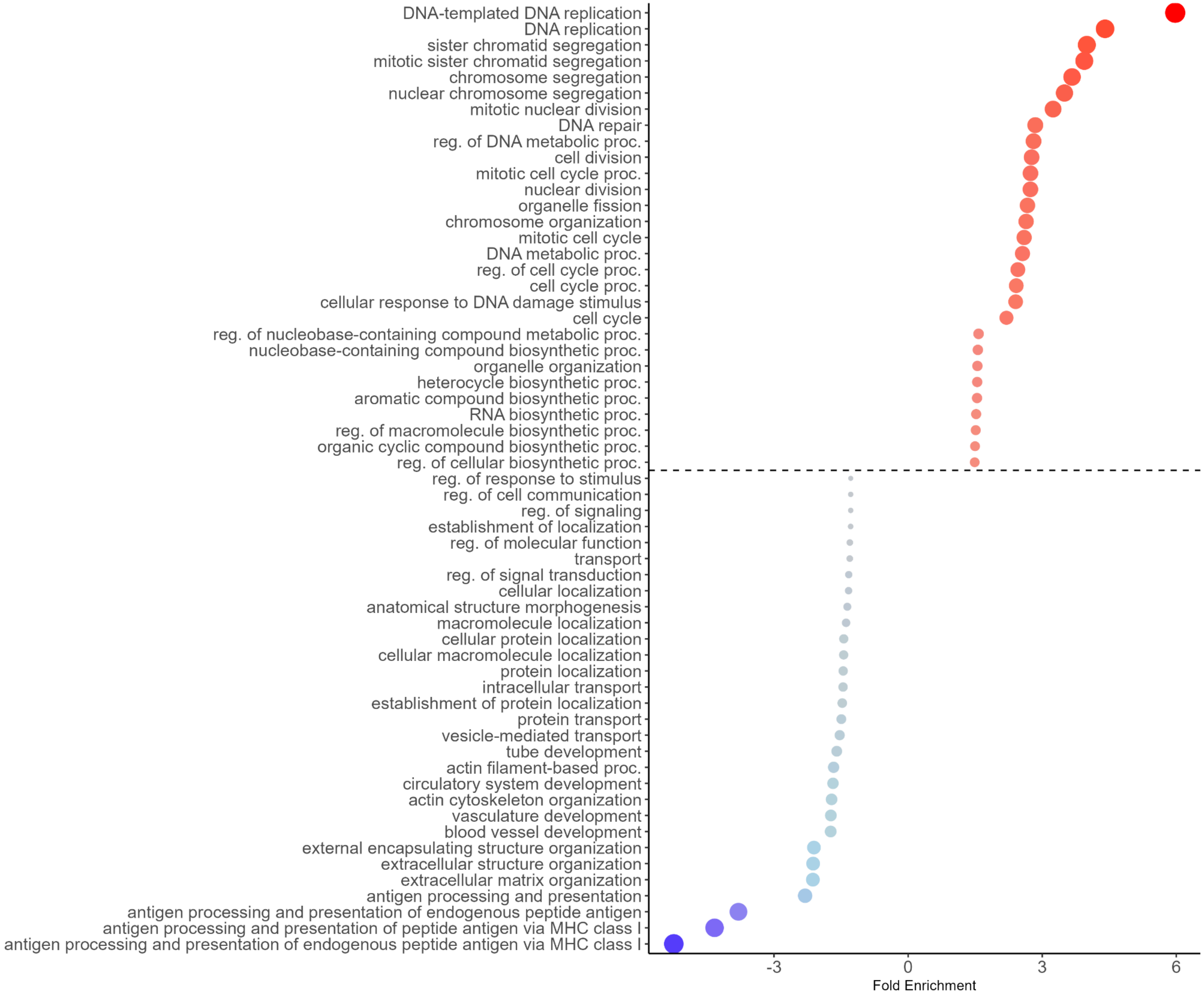


**Supplementary Figure 10.** Enrichment of biological processes for upregulated (red) and downregulated (blue) genes from cellular reprogramming (same data as Figure 2F). In this analysis we directly compared dermal fibroblasts versus reprogrammed fibroblasts.
